# A subcortical circuit linking the cerebellum to the basal ganglia engaged in vocal learning

**DOI:** 10.1101/198317

**Authors:** Ludivine Pidoux, Pascale Leblanc, Arthur Leblois

## Abstract

Speech is a complex sensorimotor skill, and vocal learning involves both the basal ganglia and the cerebellum. These subcortical structures interact indirectly through their respective loops with thalamo-cortical and brainstem networks, and directly via subcortical pathways, but the role of their interaction during sensorimotor learning remains undetermined. While songbirds and their song-dedicated basal ganglia-thalamo-cortical circuitry offer a unique opportunity to study subcortical circuits involved in vocal learning, the cerebellar contribution to avian song learning remains unknown. We demonstrate that the cerebellum provides a strong input to the song-related basal ganglia nucleus in zebra finches. Cerebellar signals are transmitted to the basal ganglia via a disynaptic connection through the thalamus and then conveyed to their cortical target and to the premotor nucleus controlling song production. Finally, cerebellar lesions impair juvenile song learning, opening new opportunities to investigate how subcortical interactions between the cerebellum and basal ganglia contribute to sensorimotor learning.

## Introduction

Speech is a highly complex motor skill which requires precise and fast coordination between vocal, facial and respiratory muscles. Human infants learn to reproduce adult vocalizations and to progressively master speech motor coordination within their first few years of life through an imitation process that builds up on motor sequence learning and strongly relies on auditory feedback (Kuhl and Meltzoff, 1996). This process, called vocal learning, is widely believed to rely on similar mechanisms as sensorimotor learning in general (Doupe and Kuhl, 1999; Kuhl and Meltzoff, 1996). The neural mechanisms underlying this process remain, however, poorly understood. Brain circuits known to be essential for sensorimotor adaptation and learning, namely the basal ganglia-thalamo-cortical loop (Krakauer and Mazzoni, 2011; Pekny et al., 2015) and the cerebello-thalamo-cortical loop (Brooks et al., 2015; Izawa et al., 2012), are both crucial for vocal learning in humans (Vargha-Khadem et al., 2005; Ziegler and Ackermann, 2017). The anatomical structure of these circuits and their function in sensorimotor learning are well conserved over vertebrate evolution (Grillner and Robertson, 2016; Redgrave et al., 1999; Sultan and Glickstein, 2007). In particular, avian song learning has been used as a paradigm to study the neural mechanisms of vocal learning, as it shares striking similarities with human speech learning (reviewed in Doupe and Kuhl, 1999).

The basal ganglia-thalamo-cortical network is involved in sensorimotor learning in several species, from lamprey to primates (Hikosaka et al., 2002; Stephenson-Jones et al., 2013; Wickens et al., 2007). The basal ganglia are thought to rely on reward prediction error signals conveyed by dopaminergic neurons (Gadagkar et al., 2016; Schultz et al., 1997; Wickens et al., 2003) to drive reinforcement learning strategies (Doya, 2000; Sutton and Barto, 1981). In songbirds, a specialized circuit homologous to the motor loop of the mammalian basal ganglia (Doupe et al., 2005) is critical for song learning in juveniles and plasticity in adults (Brainard and Doupe, 2002). This circuit is thought to correct vocal errors through reinforcement learning driven by an internal song evaluation signal conveyed by dopaminergic neurons (Fee and Goldberg, 2011; Gadagkar et al., 2016; Hoffmann et al., 2016).

The cerebello-thalamo-cortical circuit also participates in sensorimotor learning in vertebrates, from fish to primates (Brooks et al., 2015; Gómez et al., 2010; Lewis and Maler, 2004). It is believed to implement error-based supervised learning (Albus, 1971; Ito, 1984; Knudsen, 1994; Marr, 1969; Raymond et al., 1996) based on an error prediction denoting a mismatch between sensory prediction and actual sensory feedback (Doya, 2000; Dreher and Grafman, 2002). The cerebellum also drives online correction during movements building on the same sensory error prediction (Tseng et al., 2007; Booth et al., 2007). The existence of a pathway from the cerebellum to the song-related basal ganglia has been suggested by previous anatomical studies in songbirds (Person et al., 2008; Vates et al., 1997), but whether cerebellar circuits are involved in avian song learning and production remains unknown.

Beyond the indirect interaction via their respective loop with thalamo-cortical and brainstem networks, the basal ganglia and the cerebellum interact via a subcortical disynaptic pathway through the dentate nucleus, the motor part of the thalamus, and the striatum (Bostan et al., 2010; Chen et al., 2014; Hoshi et al., 2005). The cerebellum and the basal ganglia therefore do not simply act in parallel to shape cortical and brainstem activity during learning. In this paper we make the hypothesis that cerebellar signals may reach the basal ganglia to drive error correction and reinforcement learning through the same output pathway. We test this hypothesis in zebra finches. We show that (i) cerebellar inputs are conveyed via the thalamus to the basal ganglia in songbirds, (ii) they drive activity in the cortical target of the basal ganglia, and (iii) the cerebellar signals participate in juvenile song learning.

## Results

To test the hypothesis that cerebellar signals are sent to the song-related basal ganglia circuits and that the cerebellum participates in song learning, we performed the following experiments. We first reproduced the anatomical finding by Person et al. (2008) that the DCN send a projection to a thalamic region, which in turn projects to the song-related BG nucleus Area X. We then recorded responses to DCN electrical stimulation in Area X and its cortical targets and determined the nature of the neural pathway linking with pharmacological manipulations. Finally, we compared song learning ability in finches following DCN or sham lesions.

### - Anatomical connections exist from the DCN to the basal ganglia via the thalamus

We performed anatomical tracing experiments to confirm the previously reported (Person et al., 2008) indirect connection from the deep cerebellar nuclei (DCN) to the song-related basal ganglia nucleus Area X, via the dorsal thalamic zone (DTZ). The retrograde tracer Cholera-toxin B (CtB), captured by synapses (see Methods), was injected in Area X while a bidirectional tracer (fluorescently tagged dextran) was injected in the lateral DCN. Labeling of the Purkinje cells in the cerebellar cortex confirmed the proper location of the injection sites in the DCN (Fig. 1A, right panel). As shown in examples (Fig. 1B-C-D), we found fibers labeled with the DCN-injected tracer in the dorsal thalamic zone (DTZ), posterior to the thalamic nucleus involved in song learning and production (dorsolateral nucleus of the anterior thalamus, DLM), which indicates axonal projections from DCN neurons to this region. Within the same area, cell somata of thalamic neurons in DTZ were labeled with the retrograde tracer injected in Area X (Fig. 1B-C). The close association of the two types of tracers with anterogradely-labeled fibers making putative contacts on retrogradely-labeled cell bodies (Fig.1D) suggests that neurons in the lateral DCN project to DTZ thalamic neurons, which in turn project to the song-related basal ganglia nucleus Area X.

We also injected bidirectional tracers (Dextran-associated fluorochrome) in DTZ (Fig. 1E). In the cerebellum, retrograde transport of the tracer was confined to large cell bodies within the DCN (Fig. 1F). Labeled cell bodies were located for the most part in the lateral DCN. We did not find dorso-ventral distinction in the labelling in the lateral DCN, suggesting that the projection from the lateral DCN to DTZ is not topographically organized (Fig. 1F). Moreover, some neurons in the interpositus nucleus were also labeled (results not shown). This suggests that even if the projection from the cerebellum to DTZ largely comes from the lateral DCN, the interpositus nuclei may also be partially involved in this cerebello-thalamic projection. Regarding the anterograde transport of tracers injected in DTZ (Fig. 1E), we found many labeled axonal fibers in Area X, confirming the direct projection from DTZ to Area X (Fig.1G).

Thus, as already suggested in a previous study (Person et al., 2008) we found anatomical evidence for a disynaptic connection between the cerebellum and the song-related basal ganglia Area X: the lateral DCN sends projections to DTZ which in turn projects to Area X.

**Figure 1:**
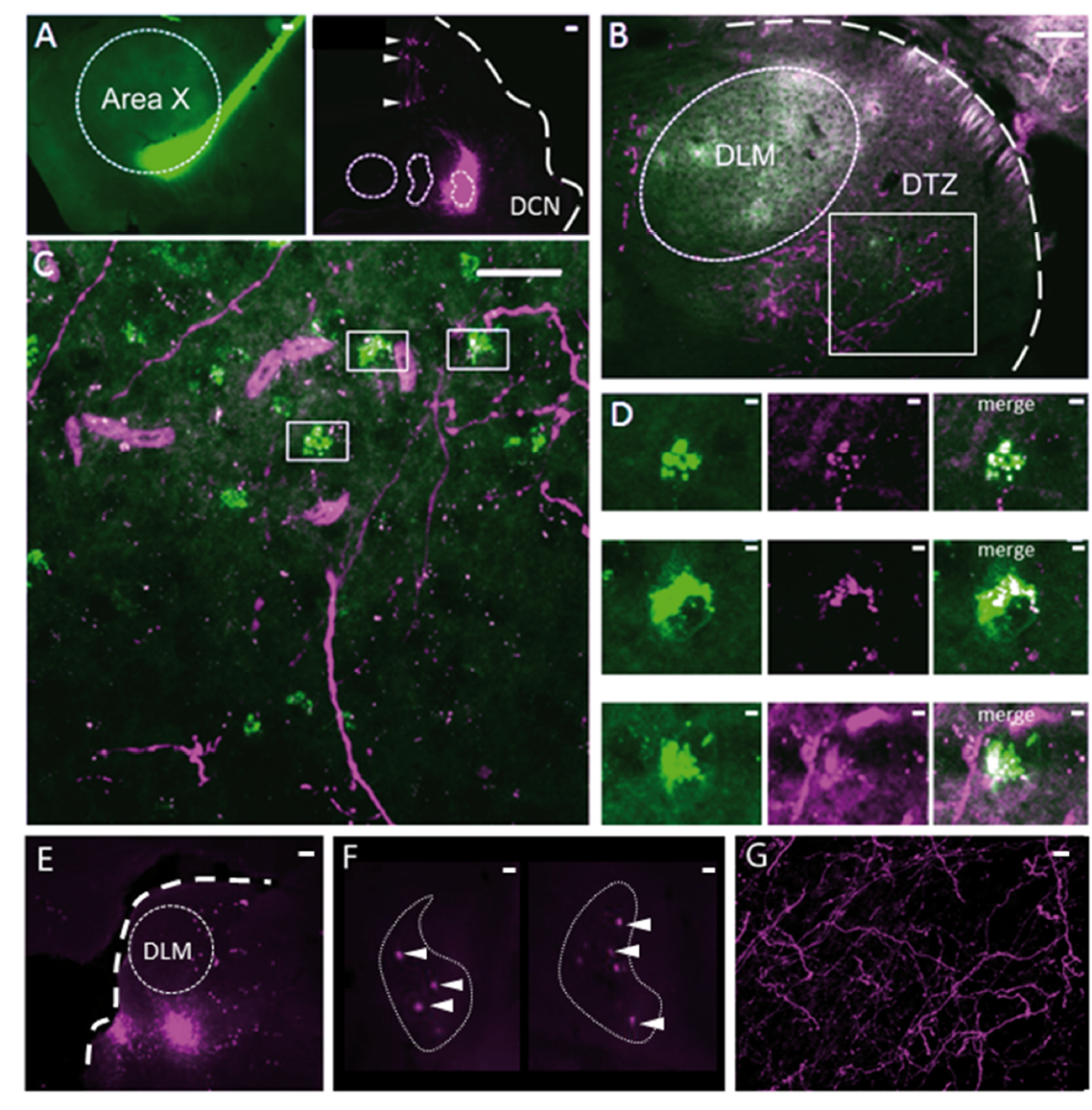
Anatomical connection between DCN and Area X. (A) Injection sites of cholera toxin B in Area X (green, left panel) and Dextran Alexa 594 in DCN (magenta, right panel). Dotted lines delimit Area X (left panel) and all three DCN (right panel). The large dotted line delimits the brain slice contour. Retrograde labeling of Purkinje cells projecting to the DCN targeted by dye injection can be observed (right panel, arrowheads). Scale bar: 100µm. (B-C): Close contacts are observed in the dorsal thalamic zone (DTZ) for a magnification of x4 (B, scale: 100µm) and x20 (C, scale: 100µm). The dotted line in B delimits nucleus DLM, while the white square in B and C indicates magnification location. Efferent fibers from Area X in DLM result in diffuse green labeling of the nucleus, while green cell somas in DTZ reflect afferent neurons. Red-labeled fibers from the DCN surround Area X-projecting neurons in DTZ. DLM: dorsolateral nucleus of the anterior thalamus. (D): Three example of close contacts between fibers from the DCN (magenta, middle panel) and soma of neurons projecting to Area X (green, left panel) in DTZ. Each panel in D corresponds to a magnification of squares indicated in C. The merge suggests an anatomical connection (right panel). Scale bar: 2µm. (E) Injection sites of Dextran Alexa 594 in DTZ. The large dotted line delimits slice contours, and the dotted circle represents DLM. Scale bar: 100µm. (F) Two examples of retrograde labelling in the lateral DCN following DTZ injection showed in E. Both examples are from the same animal, at two different depths. Arrowheads indicate DCN cell soma labelled. The dotted line delimits the lateral DCN contours. Scale bar: 20µm (G) Example of anterograde labelling in Area X. Only fibers (but no soma) are observed in Area X after DTZ injection. Scale bar: 2µm.

### -The connection from DCN to basal ganglia is functional

We then sought to determine whether the pathway revealed anatomically from the cerebellum to the basal ganglia is sufficiently efficient to drive activity within the basal ganglia. To this end we investigated the responses evoked by DCN electrical stimulation in Area X neurons.

Most neurons are silent or display very little spontaneous activity in Area X under anesthesia, whereas a minority of them displays high spontaneous activity (>25 spikes/sec, see Methods). These spontaneously active neurons are most likely pallidal-like neurons (Leblois et al., 2009; Person and Perkel, 2007). Hereafter, this population of neurons, at least some of which are area X projection neurons (Goldberg et al., 2012; Leblois et al., 2009), will be referred as pallidal neurons. DCN stimulation provoked a strong increase in the firing rate of most, if not all, pallidal neurons, as shown on the example depicted in Fig. 2B. Indeed, when a response was evoked by single-pulse stimulation in at least one pallidal neuron in Area X, all subsequently recorded neurons were also responsive to the stimulation. The response profile following DCN stimulation at a given intensity differed, however, between different pallidal neurons. This diversity of response shapes could be classified as follows: single excitatory responses (observed in 71% of case, Fig. 2C, bottom), biphasic responses with excitation followed by inhibition (observed in 19% of case, Fig.2C, middle), or triphasic responses with a rapid inhibition followed by an excitation and a late inhibition (observed in 10% of case, Fig.2C, top). A change in the response profile could also be evoked by varying the stimulation intensity: higher stimulation intensity induced biphasic or triphasic responses, while lower stimulation intensity only caused excitation. Therefore, different profiles of response can be found in the same neuron depending on the stimulation intensity used. Previous studies have shown that excitatory inputs to Area X can drive such biphasic or triphasic responses in pallidal neurons due to feedforward inhibition mediated by local inhibitory neurons (Leblois et al., 2009). The response latencies between the onset of the stimulation pulse and the onset of the excitatory response (see Methods) were broadly distributed from 10 to 50 ms (20.80 ms +/- 4.56 ms, median: 21 ms, Fig. 2D). While short latency responses (10-20 ms) can be naturally explained by a disynaptic excitatory transmission from the DCN to Area X through DTZ, biphasic and triphasic responses involve longer latencies and feedforward inhibition within Area X likely participates. Indeed, fast feedforward inhibition within Area X can delay the response of pallidal neurons to their excitatory inputs (Leblois et al., 2009), as it is the case in the mammalian striatum (Mallet et al., 2005). Altogether, these results show that stimulation of DCN neurons can drive the activity of pallidal neurons in Area X, confirming that they receive a functional input from the cerebellum.

**Figure 2:**
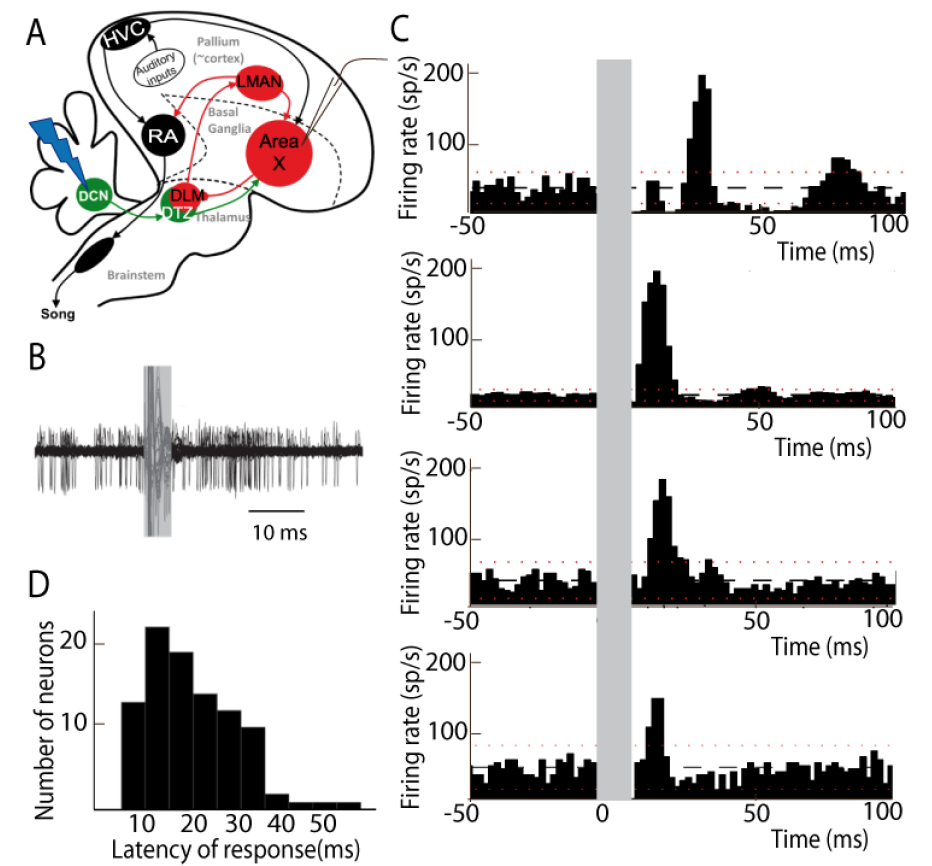
Deep cerebellar stimulation elicits strong excitation in pallidal cells of Area X. (A) Diagram of the song system in songbirds. In black, the cortical motor pathway necessary for song production. In red, the basal ganglia-thalamo-cortical loop composed of the basal ganglia nucleus Area X, the thalamic nucleus DLM, and the cortical nucleus LMAN. In green, the cerebello-thalamo-basal ganglia pathway. Stimulations are performed in the DCN during the recording of pallidal neurons in Area X. HVC: used as a proper name, RA: robust nucleus of the archopallium, LMAN: lateral magnocellular nucleus of the anterior nidopallium, DLM: medial portion of the dorsolateral nucleus of the anterior thalamus, DTZ: dorsal thalamic zone, DCN: deep cerebellar nuclei. (B) Twenty superimposed extracellular recording traces around DCN stimulation showing the increase in the number of spikes produced by a representative pallidal neuron following DCN stimulation (grey rectangle) compared to baseline firing. (C) Peri-stimulus-time-histograms (PSTHs) representing the firing rate of 4 different pallidal neurons around DCN stimulation (time bin: 2 ms). The black horizontal dashed line depicts the mean baseline firing rate and red dotted lines indicate confidence intervals (2.5 SD away from the mean baseline firing rate). Different response profiles are shown: excitation only (the two in the bottom, stimulation at 0.2 and 0.5 mA), biphasic response (second PSTH from top, stimulation at 1 mA), or inhibition and biphasic response (top, stimulation at 2 mA). (D) Distribution of response latency between DCN stimulation and the beginning of the excitatory response (20.80 ms +/- 4.56 ms, median: 21 ms).

### - The thalamic region DTZ mediates the cerebello-basal ganglia pathway

Our anatomical results strongly suggest that DTZ mediates Area X neuronal responses to cerebellar stimulation. To demonstrate that the connection is indeed functionally mediated by DTZ, we blocked glutamatergic transmission in DTZ while monitoring the responses in Area X to DCN stimulation. We pressure-injected AMPA/kainate (2,3-dihydroxy-6-nitro-7-sulfamoyl-benzo quinoxaline-2,3-dione, NBQX) and NMDA (2-amino-5-phosphonovaleric acid, APV) receptor antagonists to block all glutamatergic transmission within DTZ (see Methods, Fig. 3A) as the cerebellar projections to the thalamus are mediated by glutamate in rats (Kuramoto et al., 2009, 2011). Figure 3B shows an example of the change in the response of a pallidal neuron to DCN stimulation following the injection of glutamatergic blockers in DTZ. As our hypothesis predicts, the excitation that DCN stimulation induced in this pallidal neuron was suppressed following drug injection. We then quantified the change in response induced by glutamatergic blockers in DTZ over the population of pallidal neurons we recorded under this pharmacological protocol (n=16 pallidal neurons in 8 birds). The response strength and peak of the excitatory response (see Methods) were strongly reduced or totally suppressed when we blocked DTZ glutamatergic transmission. Mean response strength decreased from 0.55 +/- 0.13 spikes at baseline to 0.16 +/- 0.04 spikes following drug injection (paired Wilcoxon test, p=2.7642e-004, Fig.3C), and mean excitation peak from 99.35 +/- 23.41 Hz at baseline to 43.73 +/- 10.30 Hz following drug injection (paired Wilcoxon test, p=4.5523e-004). These results show that the responses to DCN stimulation in Area X pallidal neurons are mediated by glutamatergic transmission in DTZ.

Thalamo-striatal projections are glutamatergic in most vertebrates (Smith et al., 2004). It is thus natural to suppose that in zebra finches DTZ neuronal projections excite Area X neurons through glutamatergic transmission. We tested this hypothesis by blocking glutamatergic transmission around the pallidal neuron we were recording upon injection of the same drugs as above (Fig. 3D, n=8 pallidal neurons). We indeed confirmed that responses to DCN stimulation in pallidal neurons were abolished by the drug injection (Fig. 3E and F, n=8 pallidal neurons in 7 birds, response strength decreased from 0.8 +/- 0.3 spikes at baseline to 0.16 +/- 0.05 spikes following drug injection (paired Wilcoxon test, p=0.0078), and mean excitation peak from 125.01 +/- 44.19 Hz at baseline to 29.61 +/- 10.47 Hz following drug injection (paired Wilcoxon test, p=0.0078).

**Figure 3:**
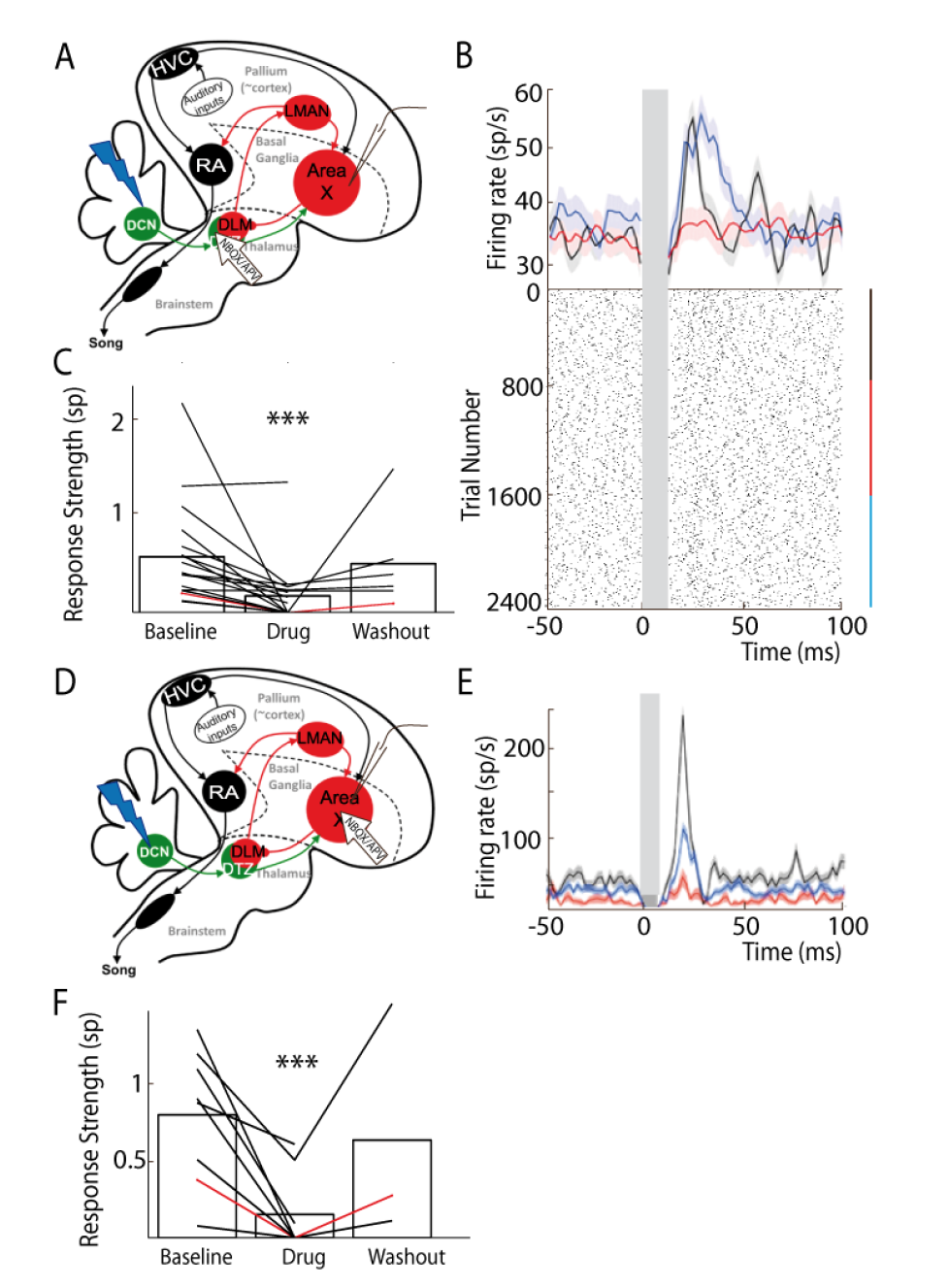
Area X pallidal responses to DCN stimulation are transmitted through excitatory synapses in DTZ and Area X. (A) Diagram of the song system in songbirds, as in Fig. 2A. Recordings are performed in Area X, NBQX/APV is applied in DTZ. (B) PSTH (top part) of a typical pallidal neuron before (black), during (red) and after (blue, washout) drug application in DTZ, and their corresponding raster plots (bottom part). (C) Population data showing the response strength of pallidal neurons in the three conditions (baseline, drug and washout, n=16 pallidal neurons in 8 birds, paired Wilcoxon test, p value<0.001). The red curve represents the example shown in B. (D) Diagram of the song system. Recordings were performed in Area X, NBQX/APV was applied in Area X in proximity to the recorded neuron. (E) PSTH representing the firing rate of one pallidal neuron, before (black), during (red) and after (blue, washout) drug application in Area X. Baseline activity after drug application (red) sometimes slightly decreases in Area X neurons compared to before drug application (black), but no significant change was observed over all neurons recorded in this condition. (F) Population data showing the evolution of response strength before, during and after drug application (n=8 pallidal neurons in 7 birds, paired Wilcoxon test, p value<0.001). The red curve represents the example shown in E.

### - LMAN does not mediate Area X responses to DCN stimulation

We cannot completely exclude that drug injected in DTZ leaked into DLM because of diffusion in the brain tissue, and that this would block a response mediated by the well-known DLM-LMAN-Area X pathway. To rule this alternative explanation out, we applied in LMAN a cocktail of AMPA and NMDA receptor antagonists while monitoring pallidal responses to DCN stimulation (Fig. 4A, top). We found no significant difference in the excitatory response of pallidal neurons to DCN stimulation between baseline and drug application conditions (Fig. 4B and 4C, n= 12 cortical neurons in 6 birds, response strength from 1.49 +/- 0.5 spikes at baseline to 1.34 +/- 0.38 spikes following drug injection, paired Wilcoxon test, p=0.479; mean excitation peak from 211.07 +/- 49.75 Hz at baseline to 200.11 +/- 47.16 Hz following drug injection, paired Wilcoxon test, p=0.5444). Following each experiment we conducted upon drug injection in LMAN, we controlled for the efficacy of the pressure injection through the glass pipette by also performing a drug injection within Area X around the recorded pallidal neurons (Fig. 4C, inset, n= 5 pallidal neurons in 5 birds, DCN stimulation response strength decreased from 1.32 +/- 0.59 spikes at baseline to 0.35 +/- 0.16 spikes following drug injection, paired Wilcoxon test, p=0.0079; mean excitation peak reduced from 182.40 +/- 81.57 Hz at baseline to 57.30 +/- 25.62 Hz following drug injection, paired Wilcoxon test, p=0.0159). These results confirm that glutamatergic transmission in LMAN is not involved in the pallidal response to DCN stimulation, ruling out a transmission through the DLM-LMAN-Area X pathway.

**Figure 4:**
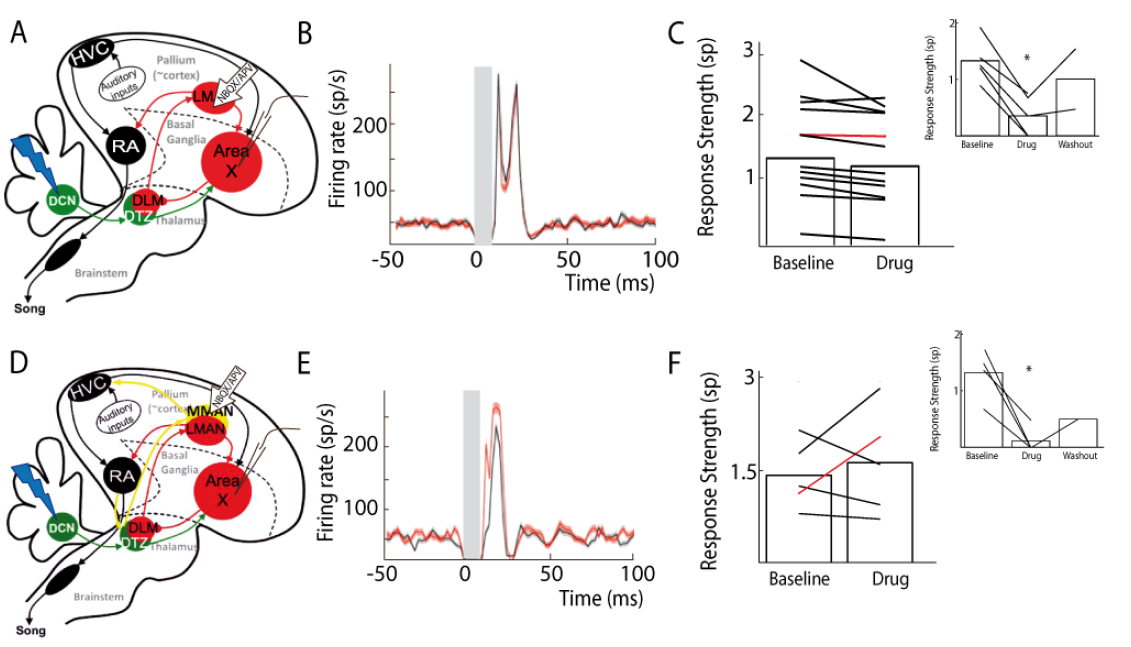
Area X pallidal responses to DCN stimulation are not transmitted through cortical nuclei LMAN or MMAN. (A) Diagram of the song system. Recordings are performed in Area X, NBQX/APV is applied in LMAN. (B) PSTH representing the firing rate of a pallidal neuron around DCN stimulation before (black) and during (red) drug application in LMAN. (C) Population data showing no change in response strength before and during LMAN glutamatergic blockade (n=12 pallidal neurons in 6 birds, paired Wilcoxon test, non-significant). The red curve represents the example shown in B. Inset: confirmation of drug efficiency by applying drug on the recorded pallidal neuron (n=5 pallidal neurons in 5 birds, paired Wilcoxon test, p<0.01). (D) Diagram of the song system. Recordings are performed in Area X, NBQX/APV is applied in MMAN, a nucleus projecting to HVC. (E) PSTH representing the firing rate of pallidal neuron before (black) and during (red) drug application in MMAN. (F) Population data showing the evolution of response strength before and during glutamatergic blockade in MMAN (n=5 pallidal neurons in 2 birds, paired Wilcoxon test, non-significant). The red curve represents the example shown in E. Inset: confirmation of drug efficiency by applying drug on the recorded pallidal neuron (n=5 pallidal neurons in 2 birds, paired Wilcoxon test, p<0.05).

### - MMAN is not involved in Area X responses to DCN stimulation

In songbirds, DTZ, receiving input from the cerebellum, is composed of several thalamic regions as described previously by anatomical studies (Person et al., 2008; Vates et al., 1997). One of these regions, called the dorsal medial posterior thalamic zone (DMP) sends a projection to the medial part of the magnocellular nucleus (MMAN) (Foster et al., 1997; Nicholson and Sober, 2015). MMAN is in turn implicated in a pathway to the song-related motor nuclei HVC (used as a proper name) and RA (Williams et al., 2012). As HVC projects to Area X in the song-related basal ganglia-thalamo-cortical circuit (Nottebohm et al., 1976, 1982), we wondered whether the response we observed in Area X could be conveyed through the MMAN-HVC-X pathway. To rule out this possibility, we blocked glutamatergic transmission in MMAN while monitoring pallidal responses to DCN stimulation (Fig. 4D). We found no significant effect of the drug injection in MMAN on the responses of pallidal neurons to DCN stimulation (Fig. 4E and 4F, n= 5 pallidal neurons in 2 birds, response strength from 1.43 +/- 0.24 spikes at baseline to 1.63 +/- 0.43 spikes following drug injection, Wilcoxon test, p=0.8125; mean excitation peak from 246.20 +/- 110.10 Hz at baseline to 258.80 +/- 115.74 Hz following drug injection, paired Wilcoxon test, p=0.4375). As previously, we checked the efficacy of the pressure injection through the glass pipette in Area X at the end of each experiment (Fig. 4F, inset, n= 5 pallidal neurons, from 1.33 +/- 0.66 spikes at baseline to 0.12 +/- 0.06 spikes following drug injection, Wilcoxon test, p=0.0286; mean excitation peak from 238.50 +/- 119.25 Hz at baseline to 35.00 +/- 17.50 Hz following drug injection, paired Wilcoxon test, p=0.0268). This experiment ruled out the possible transmission of pallidal responses to DCN stimulation through the MMAN-HVC-Area X pathway.

In conclusion, the results of our electrophysiological experiments provide strong evidence that the cerebellum is linked to the song-related basal ganglia nucleus Area X through a functional excitatory connection involving a glutamatergic projection from the DCN to DTZ, and a glutamatergic projection from DTZ to Area X.

### - The cerebellar responses are conveyed to LMAN through the basal ganglia loop

In songbirds, Area X is known to be part of the basal ganglia-thalamo-cortical circuit homologous to the motor loop of the basal ganglia-thalamo-cortical networks in mammals (Brainard and Doupe, 2002). In the following experiments we tested whether responses observed in the pallidal neurons after DCN stimulation were conveyed to the output nucleus of the basal ganglia-thalamo-cortical loop, namely LMAN (Fig.5A). We recorded LMAN neurons and found that DCN stimulation elicited strong responses in all LMAN neurons recorded (Fig. 5B). This response is composed of two excitatory components: a strong and rapid excitation, and a long and slow one. Such bimodal excitatory response with two peaks was found in 10% (n=3/30) of the LMAN neurons recorded. For the majority of recorded LMAN neurons (90%, n=27/30), we saw only one of the two excitatory phases provoked by DCN stimulation. The latency of excitatory responses in LMAN neurons was therefore spread in a bimodal distribution (Fig. 5C) with two widely different peaks: a first peak between 10 and 50 ms (26 +/- 7.8 ms, median: 19ms, 28% of all recorded LMAN neurons, n=8/30), and a second peak around 100 ms (125 +/- 32 ms, median: 110 ms, 72 % of all recorded LMAN neurons, n=22/30). Interestingly, these two peaks in the latency distribution in LMAN neurons mirrored the inhibitory responses observed in Area X pallidal neurons. Indeed, Area X neurons displayed inhibitory responses either preceding or following the excitatory component of their response. An inhibition in Area X pallidal neurons, many of which project to the thalamic nucleus DLM (Fee and Goldberg, 2011; Leblois et al., 2009), induces a fast excitatory response in DLM neurons (Goldberg et al., 2012; Leblois et al., 2009; Person and Perkel, 2007) and thereby activates LMAN through DLM excitatory projections (Leblois et al., 2009). The first excitation in LMAN neurons, around 20 ms latency, could therefore be mediated by the fast inhibition observed in pallidal neurons (Fig.2C, top panel). Similarly, the slow inhibitory component of pallidal responses to DCN stimulation, with a mean latency of 28ms (28.2 +/- 9.5 ms, data not shown), likely activates the DLM-LMAN pathway with much longer latencies (>50 ms) and may therefore drive the second excitation in LMAN. To confirm that the response in LMAN neurons is mediated by Area X, we blocked glutamatergic transmission in Area X to prevent local responses to DCN stimulation (Fig.5D, left panel). Hereafter, the response strength is calculated as the total area of the response, containing the two peaks of excitation when they are present. After application of the glutamatergic blockers to Area X, responses disappeared in LMAN (Fig.5E), with a significant reduction or suppression of excitatory response over all LMAN neurons recorded (Fig.5F, n= 14 multiunit recording, from 2.04 +/- 0.54 spikes at baseline to 0.89 +/- 0.23 spikes following drug injection, paired Wilcoxon test, p=0.0012; mean peak excitation from 27.31 +/- 7.29 Hz at baseline to 10.88 +/- 2.90 Hz following drug injection, paired Wilcoxon test, p= 3.6621e-004).

**Figure 5:**
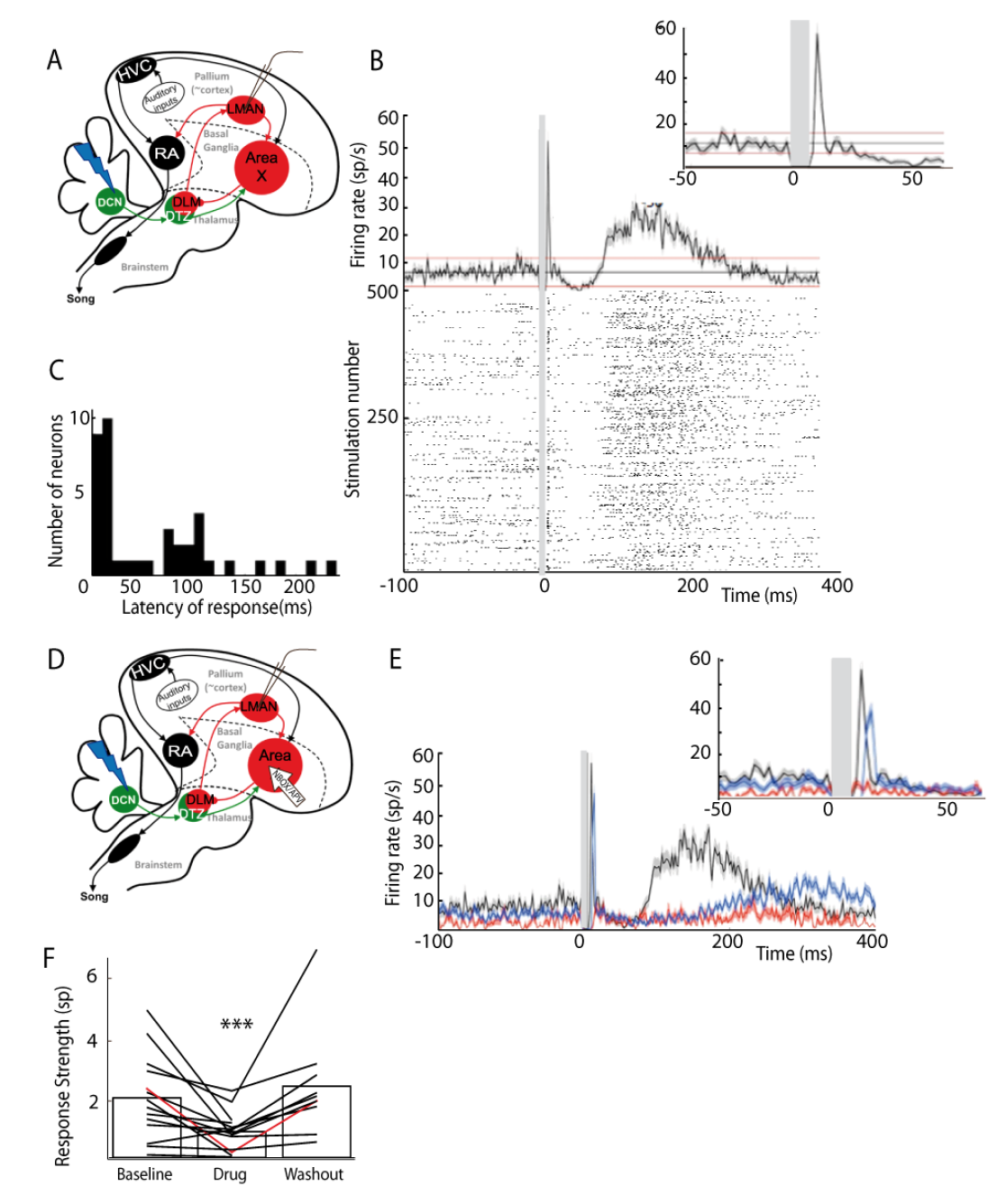
LMAN neurons display bimodal responses to DCN stimulation mediated by Area X. (A) Diagram of the song system, as in Fig. 2A. Neurons were recorded in LMAN during DCN stimulation (B) Example response in a typical LMAN recording following DCN stimulation with the corresponding raster plot (multiunit recording). Inset: magnification of the first excitatory peak. (C) Distribution of response latency over all LMAN recordings displaying the two characteristic peaks of response (first peak: 10-30 ms and second peak: 100 ms, see Results, time bin: 10ms). (D) Diagram of the song system, as in Fig. 2A. NBQX/APV is applied in Area X and neurons are recorded in LMAN during DCN stimulation. (E) Example response following DCN stimulation from a typical recording in LMAN (multiunit recording), before (black), during (red) and after (blue, washout) the drug application. (F) Population data showing the evolution of response strength over the three periods (baseline, drug, washout, n= 14 multiunit recording in 5 birds, paired-Wilcoxon test, p-value=0.001). The red curve represents the example shown in E.

### - The cerebellum can influence the discharge of RA neurons

The basal ganglia-thalamo-cortical loop affects song production and drives song plasticity via its projection to nucleus RA (Andalman and Fee, 2009; Bottjer et al., 1984). We tested whether DCN stimulation also drives responses in RA neurons via the basal ganglia-thalamo-cortical loop (Fig. 6A). DCN stimulation induced strong excitatory responses in RA neurons (Fig. 6C, black curve) with latencies in the range from 10 to 100 ms (30.2ms +/- 7.8 ms, median: 16 ms). Blocking glutamatergic transmission in LMAN significantly reduced the excitatory response to DCN stimulation in RA neurons (Fig. 6C and 6D, n=6 neurons in 5 birds, response strength decreased from 0.8 +/- 0.32 spikes at baseline to 0.29 +/- 0.12 spikes following drug injection, Wilcoxon test, p =0.0087; mean excitation peak from 186.66 +/- 76.20 Hz at baseline to 71.18 +/- 29.06 Hz following drug injection, paired Wilcoxon test, p= 0.0156).

**Figure 6:**
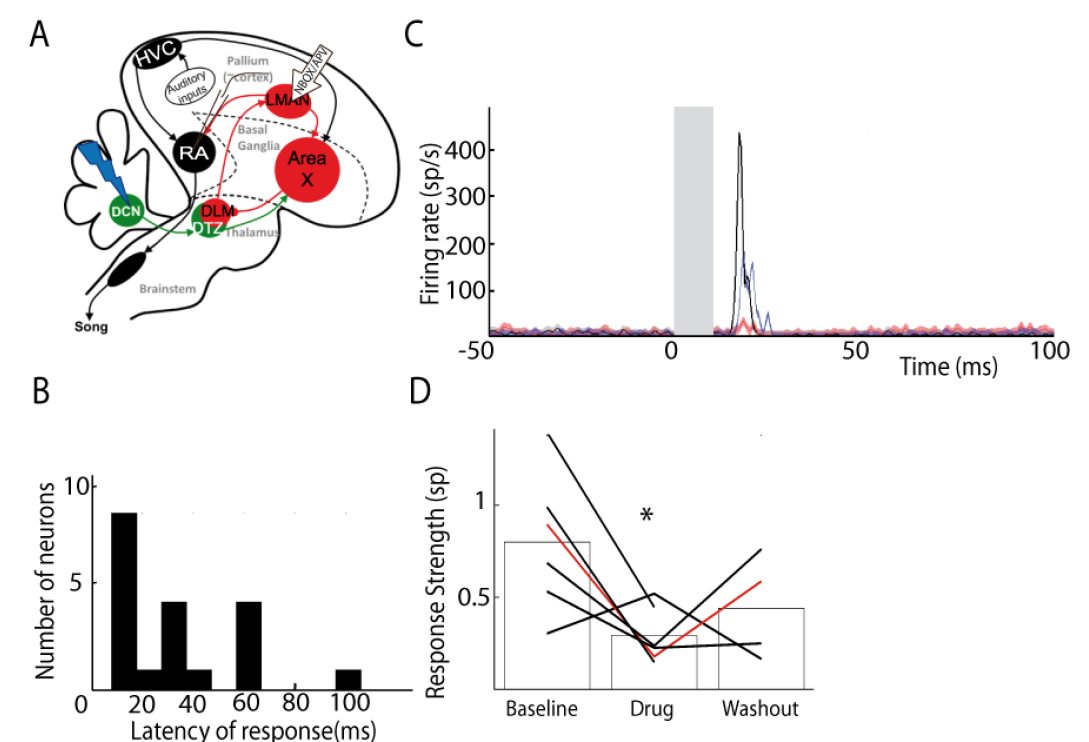
RA responses to DCN stimulation can be partially suppressed by blocking glutamatergic transmission in LMAN. (A) Diagram of the song system, as in Fig. 2A. Neurons were recorded in RA during DCN stimulation, NBQX/APV was applied in LMAN. (B) Distribution of RA neurons response latencies (time bin: 10ms). (C) PSTH representing the firing rate of a typical RA neuron before (black), during (red) and after (washout, blue). (D) Population data showing the change of response strength over the three periods (baseline,drug, washout, n=6 neurons in 5 birds, paired Wilcoxon test, p-value<0.05). The red curve represents the example shown in C.

### - DCN lesion impairs song learning in juvenile zebra finches

Our experiments provide strong evidence for a functional disynaptic cerebellum-thalamus-basal ganglia pathway in songbirds. This pathway can drive the output nucleus of the basal gangliathalamo-cortical loop, LMAN, and can influence the premotor activity of RA neurons.

Song learning strongly relies on the basal ganglia-thalamo-cortical loop (Bottjer et al., 1984; Brainard and Doupe, 2002; Nottebohm et al., 1976; Scharff and Nottebohm, 1991). As we demonstrated that the cerebellar connection to the basal ganglia-thalamo-cortical loop is efficient, we tested the hypothesis that the cerebellum contributes to song learning. Juvenile zebra finches were subjected to partial lesions in their lateral DCN, either electrolytic (n=7) or chemical using ibotenic acid (n=3). Figure 7D displays the spectrograms of the song motifs produced by a tutor and its two fledglings, one of them with a DCN lesion. The juvenile bird that underwent the DCN lesion copied fewer syllables than his control brother. We compared the quality of tutor imitation in young male juveniles undergoing partial DCN lesion or sham surgery. To this end, we compute a similarity score based on the peak crosscorrelation between the spectra of the tutor’s motifs and of the juvenile’s songs. This score may be affected by both acoustic and temporal features of the song (see methods). In sham-lesion juvenile birds, the similarity score between the juvenile’s song and the tutor’s song motif was increased between the day preceding the sham-lesion surgery and the crystallization stage (before lesion: 0.27 ± 0.12; at 90 dph: 0.44 ± 0.09; paired Wilcoxon test, p=0.04, df=5; Fig 7E). On the contrary, tutor song imitation did not improve in juvenile birds following partial lesions in the lateral DCN (before lesion: 0.33 ± 0.15; at 90 dph: 0.35 ± 0.13; paired Wilcoxon test, p=0.06, df=9; Fig 7E). Moreover, there was a significant correlation between the proportion of the lateral DCN that was left unaffected and the quality of the tutor song imitation (r=0.57, p=0.03; Fig 7F). In adult birds, DCN lesion did not induce any detectable change in syllable acoustic features (results not shown). In conclusion, our lesion experiment shows that the cerebellum contributes to song learning in juvenile zebra finches.

**Figure 7:**
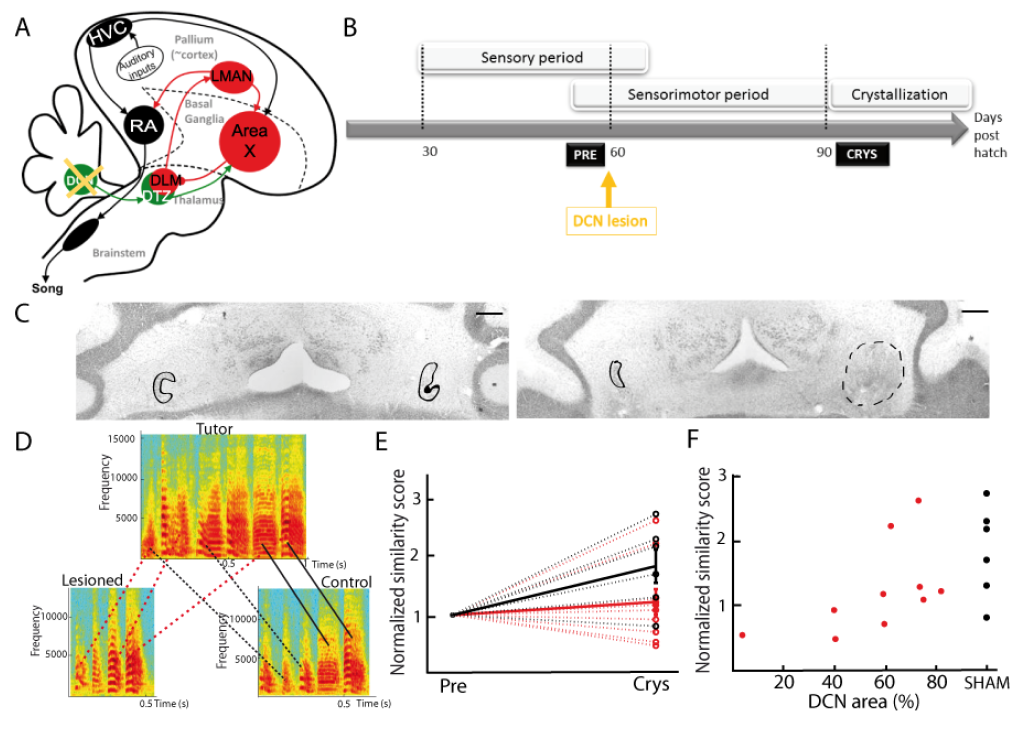
DCN lesions impair song learning and reduce the similarity to tutor song after crystallization. (A) Diagram of the song system, as in Fig. 2A, representing DCN lesion. (B) Diagram of the song learning periods in songbirds: the sensory period, the sensorimotor period in which juveniles start to produce sounds, and the crystallization phase. Lesions were made at 60 dph. (C) Nissl staining on horizontal slices showing the deep cerebellar nuclei. The black lines delimit in the two hemispheres the lateral nuclei. The dotted line (right panel) delimits the lesion site. Left: control bird. Right: bird with DCN lesion. (D) Examples of three spectrograms of tutor and juveniles song motifs at crystallization: top: tutor song motif, bottom left: song motif of a juvenile with DCN lesion, bottom right: control juvenile. Solid lines connect two similar syllables found in the tutor and juvenile song motifs, dotted lines between two syllables reflect a partial copy of the tutor syllable (red lines for the juvenile with DCN lesion, black lines for the control juvenile). (E) Population data showing the evolution of similarity between the day before the lesion (pre) and the crystallization period (90 dph) in juveniles with sham lesions (black dots for individuals, solid black line for the mean) and DCN lesions (red dots for individual, solid red line for the mean). Data are normalized over the pre-lesion period (see Methods for the normalization, n=10 birds with lesion, n=6 sham birds, Wilcoxon test, p<0.05). (F) Normalized similarity score plotted as a function of the total area left from the lateral DCN (%) for juveniles with DCN lesion (red dots, n=10 lesion birds) or sham lesion (black dots, n=6 sham birds). A significant correlation was revealed between the similarity and the proportion of lateral DCN left intact (r=0.57, p<0.05).

## Discussion

Previous investigations into the neural mechanisms of vocal learning in songbirds have focused on the contribution of pallial and basal ganglia circuits (Mooney, 2009), ignoring a possible contribution of the cerebellum to avian song learning. Here, we bring strong evidence that the cerebellum interacts with song-specific circuits in the basal ganglia and participates to song learning in juvenile birds. Indeed, we have demonstrated that the DCN project via a disynaptic pathway to the song-related basal ganglia nucleus Area X and that the cerebellum is able to modulate Area X output, its cortical target, and the premotor nucleus RA, via a thalamic relay. These results are reminiscent of the cerebello-thalamo-basal ganglia pathway recently discovered in mammals (Bostan et al., 2010; Chen et al., 2014). We also demonstrate that the cerebellum contributes to song learning, as a lesion in the DCN impaired song learning in juvenile birds.

### - Partial lesions in the cerebellum

The DCN receive strong convergent input from the inferior olive and from Purkinje cells from many functional territories in the cerebellar cortex (Apps and Garwicz, 2005). Given this strong convergence of multi-modal inputs to the DCN, large bilateral lesions in the DCN can strongly impair vital sensorimotor abilities potentially leading to a high post-operative mortality. We therefore limited the extent of our lesions to reduce the impact on global function. Still, transient motor impairments were observed during first couple of days following surgery that disappeared rapidly as the birds resumed perching and singing. While behavioral monitoring ensured that global functions were normal when we quantified the effect of lesions on song, we cannot totally exclude the fact that non-specific motor effects of the lesions were partially responsible for our song-specific behavioral results. To rule out this experimental limitation regarding partial lesions, specific lesions of the cerebellothalamic projection should be performed in the future.

### - Several types of Area X neuron are potentially involved in the cerebello-thalamo-basal ganglia pathway

Our results indicate that the cerebellar input to the basal ganglia modulates the activity of pallidal neurons in Area X, but we did not directly investigate the response of other neuronal types in this structure. Area X contains all the neuron types found in the striatum and pallidum in mammals (Farries and Perkel, 2000, 2002): pallidal neurons, medium spiny neurons and many striatal interneuron types. Only pallidal neurons, however, project outside of Area X; these share physiological, biochemical and anatomical properties of mammalian pallidal neurons (Carrillo and Doupe, 2004). Area X pallidal neurons display strong spontaneous activity both *in vitro* (Budzillo et al., 2017; Farries and Perkel, 2000, 2002) and *in vivo* (Person and Perkel, 2007; Goldberg and Fee, 2010) and can therefore be distinguished from the other neuronal populations in Area X, the spontaneous activity of which is much lower (Person and Perkel, 2007; Leblois et al., 2009; Goldberg and Fee, 2010). Given the strongly bimodal distribution of spontaneous activity observed in our recording (see methods) and the relative scarcity of neurons displaying a low spontaneous activity in Area X (Farries and Perkel, 2002), our dataset is likely to contain mostly if not only pallidal neurons. A contribution from a small fraction of spontaneous striatal interneurons cannot, however, be ruled out.

### - Similarities and differences between the cerebello-thalamo-basal ganglia pathways of mammals and songbirds

In mammals, a pathway connecting the cerebellum and the striatum through the thalamus was demonstrated in rodents (Chen et al., 2014) and monkeys (Hoshi et al., 2005). However, it remains unknown whether and how these cerebellar inputs are conveyed to basal ganglia output neurons and to their thalamo-cortical targets ultimately affecting behavior (Alexander, 1994; Alexander et al., 1990). Here, we show in songbirds that the cerebellar signals travel through the basal ganglia-thalamo-cortical circuit and can drive firing in song-related premotor neurons in RA. In monkeys, the dentate nucleus can be divided into two parts: the dorsal part, which has reciprocal projections with motor and premotor cortical areas via the motor thalamus, and the ventral part, which has reciprocal projections with associative and other non-motor cortical areas via non-motor thalamic regions (Dum and Strick, 2003; Kelly and Strick, 2003; Orioli and Strick, 1989). Additionally, anatomical tracing showed that some projections to the thalamus also come from the interpositus and the fastigial nuclei (25%) (Bostan et al., 2010; Hoshi et al., 2005). In songbirds, our tracing experiments showed that DTZ projects to the song-related basal ganglia nucleus Area X and receives extensive axonal projections from the most lateral of DCN, analogous to the dentate nucleus in mammals (Arends and Zeigler, 1991; Sultan and Glickstein, 2007; Voogd and Glickstein, 1998). We found no dorso-ventral contrast in our anatomical results and thus make no distinction between potential motor and non-motor parts of the lateral nucleus. Bidirectional tracer injections in DTZ however revealed a weaker, but consistent, projection from the intermediate nucleus, analogous to nucleus interpositus in mammals (Arends and Zeigler, 1991; Sultan and Glickstein, 2007; Voogd and Glickstein, 1998). The anatomical labeling in the intermediate nucleus was less intense compared to the lateral nucleus, suggesting that it sends weaker projections to the thalamus. Both nuclei seem, however, to project to DTZ and may thereby be involved in the cerebello-thalamo-basal ganglia pathway studied here.

During our electrophysiological experiments, the stimulation electrode placement targeted the most lateral DCN, as confirmed histologically. Although unlikely, we cannot completely exclude that the stimulation current could spread into the intermediate nucleus and activate projection neurons there as well. Indeed, the size of the stimulated area is not well controlled (Ranck, 1975; Tehovnik et al., 2006). This raises the possibility that the intermediate nucleus could also be involved in the neural responses observed in the basal ganglia-thalamo-cortical loop following DCN stimulation. Further investigations to assess the role of the putative connections between the intermediate nucleus and the thalamus therefore remain needed.

In monkeys, the dorsal part of striatum, dedicated to motor functions, receives thalamic projections (Parent and Hazrati, 1995). In songbirds, the song-related basal ganglia nucleus Area X is a rostro-ventral structure. The ventral position of this nucleus is an unusual feature of the song system given its motor function (Brainard and Doupe, 2002). Because Area X contains a mixture of striatal and pallidal neurons (Farries and Perkel, 2000, 2002), we were not able to distinguish if the thalamic fibers arrive on striatal neurons firstly as it has been shown in mammals (Smith et al., 2004), on the pallidal neurons directly, or both. While we focused on the thalamic projection on Area X, thalamic projections may also reach other parts of the avian basal ganglia. Determining which thalamic area projects to which neurons in the basal ganglia will however require multiple tracing studies and was therefore left for future investigations.

### - Is the cerebello-thalamo-basal ganglia pathway the only functional pathway connecting cerebellum to the song system?

Beyond a subcortical connection between the dentate nucleus and the basal ganglia described in mammals, the cerebellum is also known to project to the motor part of the thalamus, which in turn projects to the motor cortex (Kelly and Strick, 2003). This disynaptic connection between the cerebellum and the motor cortex is known to be important in motor control and motor coordination (Brooks, 1984). In songbirds, RA is considered as a premotor nucleus (McCasland, 1987), and its efferent projections are equivalent to descending projections from M1 to brainstem and spinal circuits in mammals (Medina and Reiner, 2000; Zeier and Karten, 1971). While we have revealed a subcortical connection between the cerebellum and basal ganglia which indirectly affects premotor activity in RA, no direct connection from the cerebellum to a thalamo-cortical circuit including RA has been described yet in songbirds. Nevertheless, DTZ, which mediates cerebellar input to the basal ganglia, is also known to project to the pallial nucleus MMAN, which in turn projects to HVC (Foster et al., 1997; Nicholson and Sober, 2015; Williams et al., 2012). As HVC directly projects to RA through the cortical pathway controlling song production, a DCN-DTZ-MMAN-HVC-RA projection may represent the functional equivalent of the mammalian cerebello-thalamo-cortical pathways. This new pathway should be characterized by anatomical and electrophysiological experiments to assess the impact of cerebellar input on the cortical pathway during song learning and production.

### - Potential impact of cerebellar input on basal ganglia

We have shown that a cerebello-thalamo-basal ganglia pathway exists in songbirds, is functional and shares many similarities with the mammalian cerebello-thalamo-basal ganglia pathway. Knowing the role of the cerebellum and the basal ganglia respectively in supervised and reinforcement learning (Doya, 2000), we hypothesize that the cerebellum can participate in basal ganglia functions by sending an error-correction signal related to a detected mismatch between actual and predicted sensory feedbacks. This error correction signal will be integrated into the basal ganglia to drive the motor command output during the learning process. In this hypothesis, both the reward prediction error signal driving reinforcement learning and the cerebellar error correction signal would cooperate within the basal ganglia to achieve faster and more efficient sensorimotor learning. Alternatively, the cerebellar input could modulate the cortico-striatal plasticity (Chen et al., 2014) and thereby regulate the learning rate in basal ganglia circuits.

In songbirds, motor variability and error correction in song involve the basal gangliathalamo-cortical loop (Andalman and Fee, 2009; Kao and Brainard, 2006; Olveczky et al., 2005; Tumer and Brainard, 2007). Indeed, this circuit is necessary for the induction of song plasticity (Andalman and Fee, 2009; Brainard and Doupe, 2002). Lesions (Olveczky et al., 2005; Tumer and Brainard, 2007) or reversible inactivation (Andalman and Fee, 2009) of the output of the cortico-basal ganglia loop reduces song variability and impairs error correction during song learning (Andalman and Fee, 2009; Olveczky et al., 2005; Tchernichovski et al., 2001). These functions presently attributed to the basal ganglia-thalamo-cortical loop could also be influenced by the cerebellum through its subcortical connection to the basal ganglia nucleus Area X.

The cerebellum is implicated in diverse sensorimotor processes (Ackermann, 2008; Izawa et al., 2012) and cerebellar lesions prevent good performance in sensorimotor tasks like reaching (Izawa et al., 2012), vocal production (Ackermann, 2008) or the vestibulo-ocular reflex (Ito, 1998), among others. Moreover, the subcortical pathway from cerebellum to basal ganglia is involved in dystonia (Calderon et al., 2011; Fremont et al., 2017; Neychev et al., 2008; Tewari et al., 2017). The existence of the cerebello-thalamo-basal ganglia pathway makes the songbird model, classically used as a model to study vocal learning, a good model for further investigations of the cooperation between cerebellum and basal ganglia in sensorimotor learning and its dysfunction in movement disorders.

## Materials and Methods

### - Animals

All the experiments were performed in adult male zebra finches *(Taeniopygia guttata*), >90 days post-hatch unless otherwise specified. Birds were either reared in our breeding facility or provided by a local supplier (Oisellerie du Temple, L’Isle d’Abeau, France). All animals had constant access to seeds, crushed oyster shells and water. Seeds supplemented with fresh food and water were provided daily. Birds were housed on a natural photoperiod (both in the aviary and in sound isolation boxes during the behavioral experiment). Animal care and experiments were carried out in accordance with the European directives (2010-63-UE) and the French guidelines (project 02260.01, Ministère de l’Agriculture et de la Forêt). Experiments were approved by *Paris Descartes University* ethics committee (Permit Number: 13-092).

### - Surgery

Before surgery, birds were first food-deprived for 20-30 min, and an analgesic was administered just before starting the surgery (meloxicam, 5 mg/kg). The anesthesia was then induced with a mixture of oxygen and 3-5% isoflurane during 5 minutes. Birds were then moved to the stereotaxic apparatus and maintained under anesthesia with 1% isoflurane. Xylocaine (31.33mg/mL) was applied under the skin before opening the scalp. Small craniotomies were made above the midline reference point, the bifurcation of the midsagittal sinus, and above the structures of interest. Stereotaxic zero in anteroposterior and mediolateral axis was determined by the sinus junction. To ease the access to the cerebellum, we used a head angle of 50°. The stereotaxic coordinates used for each brain structure are summed up in Table 1.

**Table 1:** Stereotaxic coordinates summary. Head and arm angle (on the mediolateral axis) are expressed in degrees, anteroposterior and mediolateral coordinates are expressed in millimeters from the sinus junction, and depth coordinates in millimeters from the surface of the brain. DCN: deep cerebellar nuclei. LMAN: lateral magnocellular nucleus of the nidopallium, MMAN: medial magnocellular nucleus of the nidopallium, HVC: used as a proper name, DTZ: dorsal thalamic zone.

| Structure | Head angle (°) | Arm angle (°) | Antero-post (mm) | Medio-lateral (mm) | Depth (mm) |
| --- | --- | --- | --- | --- | --- |
| Area X | 50 | 0 | 4.0 | 1.5 | 3.0-4.0 |
|  | 50 | 15 | 4.0 | 2.7 | 3.5-4.5 |
| DCN | 50 | 15 | -2 | 2.5 | 3.5 |
|  | 50 | 0 | -1.5/-1.8/-2.1 | 1.3 | 3.4 |
| DTZ | 50 | 0 | 0-0.3 | 1.2 | 4.3-4.5 |
| LMAN | 50 | 0 | 4.1 | 1.8 | 2.3-2.5 |
|  | 50 | 15 | 4.1 | 3.0 | 2.4-2.6 |
| MMAN | 50 | 0 | 4.1 | 0.5 | 2.3-2.5 |
|  | 50 | 15 | 4.1 | 1.7 | 2.4-2.6 |

### Anatomical tracing

We performed iontophoretic injections of fluorescent dye using dextran conjugates with Alexa 594 (Thermofischer, 5% in PBS 0.1M 0.9% saline) in targeted cerebral structures (lateral DCN and Area X nucleus) using a glass pipette with a small (10µm) tip and ±5 µA DC pulses of 10 s duration, 50% duty cycle, applied for 3 min. In the cerebellum, to be sure that the injection was constrained to the lateral deep cerebellar nucleus, we verified that the retrograde labeling of Purkinje cells was limited the most lateral sagittal zone (Fig.1A).

In additional tracing experiments, 250 nL of cholera toxin tracers coupled with Alexa 488 (Thermofischer, diluted in PBS 0.1M 0.9% saline) were pressure-injected with a Hamilton syringe (1µL, Phymep, Paris, France), at 100nL per minute, at each injection site (2 injection sites per brain hemisphere). Birds were then housed individually for three days after injection to allow for dye transport.

### - In vivo electrophysiology

Recordings in Area X, LMAN, and RA were made with a tungsten electrode with epoxy isolation (FHC, impedance varying from 3.0 to 8.0 MΩ depending on the type of neuron recorded). Acquisition of the signal was done with the AlphaOmega software, using low-pass (frequencies below 8036 Hz) and high-pass (frequencies above 268 Hz) filters to only detect the spike signal. The sampling frequency was 22320 Hz. In area X, the recorded neurons displayed a bimodal distribution of spontaneous firing rate, above 25 Hz or under 10 Hz. We considered neurons with frequency above 25Hz as pallidal neurons in Area X (Leblois et al., 2009; Person and Perkel, 2007). Others neurons in Area X with spontaneous firing rates under 10Hz were not taken into account in the present study. Note that the level of spontaneous activity is different under anesthesia compared to what was seen in awake birds (Goldberg et al., 2010) and can vary depending on the specific drug used (Brooks, 1984). This may explain the slight difference in spontaneous activity among neurons recorded here as pallidal, compared to previous studies performed under urethane anesthesia (Leblois et al., 2009; Person et al., 2007), known to preserve awake-like cortical activity (Albrecht et al., 1990).

A single-pulse electrical stimulation in the lateral deep cerebellar nucleus (DCN) was applied through a bipolar electrode during recording of different structures in the contralateral basal ganglia nucleus (Area X), the lateral part of the magnocellular nucleus (LMAN), the medial part of the magnocellular nucleus (MMAN), and robust archopallium nucleus (RA). The duration of the stimulation was 1 ms, with an inter-stimulation time of 1.6 s, and the intensity ranged from 0.1 to 4 mA. Despite long stimulation duration, observed responses in recorded neurons were stable over time. We aimed to place the stimulation electrode within the lateral cerebellar nucleus, and the positioning of the electrode was confirmed histologically (see next paragraph). However, we cannot completely rule out that the stimulation current did spread to the nearby interpositus nucleus.

### - Pharmacology

During electrophysiological experiments, drugs were applied locally by pressure with small tip glass pipette (10µm) and nitrogen picospritzer (Phymep, Paris, France). We used a mix of NBQX 5mM (Sigma Aldrich, diluted in PBS 0.1M 0.9% saline) and APV 1mM (Sigma Aldrich, diluted in PBS 0.1M 0.9% saline) to block glutamate receptors.

### - Data analysis

Analyses of recorded neurons after DCN stimulation were done using Spike 2 and Matlab. Spike sorting was performed with the software Spike2 (CED, UK), using principal components analysis of spike waveforms. For Area X neurons, and RA neurons, we managed to record single units, and we focus on these single unit neurons in the analysis. In the LMAN and MMAN, we chose to record mostly multiunit activity. Indeed, most neurons in these nuclei exhibit very low spontaneous activity (∼1 sp/s), leading to wide fluctuation in the PSTH estimate of baseline activity preceding stimulation with high temporal resolution (time bin: 10ms) and making it difficult to estimate response latency, strength and duration. Instead multi-unit activity with higher baseline levels allows better baseline statistics and narrower confidence intervals for the detection of the response to stimulation.

Spike train analysis was then performed using Matlab (MathWorks, Natick, MA, USA). We calculated peri-stimulus time histograms (PSTH) of recorded neurons after DCN stimulation. PSTHs were calculated with a 2-ms bin for neurons in Area X and RA. For structures with low firing rate (LMAN and MMAN) the time bin was 10 ms to limit bin-to-bin fluctuations in spike count. We calculated the mean and the standard deviation (SD) of the firing rate over the period preceding the stimulation (50ms for Area X and RA, 100 ms for LMAN and MMAN), and we considered that a neuron exhibited a significant response to the stimulation when at least two consecutive bins of the PSTH were above (for excitation) or below (for inhibition) the spontaneous mean firing rate +/- 2.5*SD. The return of two consecutive bins at the spontaneous mean firing rate +/- 2.5*SD indicated the end of the response. We defined the latency of response as the time between the stimulation onset and the beginning of the first excitatory or inhibitory response. Response strength was calculated as the sum of the difference between the PSTH values and the mean baseline firing rate over the entire response period, and represents the average number of excess (default) spikes induced by a single stimulation. For neurons in Area X and RA, the response strength was calculated over the first peak of excitation only (as most responses did not elicit two peaks of excitation, see Results). For LMAN and MMAN neurons recording, neurons tended to display bimodal responses (see Results) and both the first and second excitation peaks were taken into account to calculate the response strength. We also report the peak firing rate in the response period as the maximal value of the PSTH. The PSTHs are displayed either as histograms or as solid curves with gray shaded area surrounding the curve representing the SD of the baseline firing rate.

### - Lesion experiments

Lesions were performed in the DCN of juvenile zebra finches. We targeted the most lateral DCN, analogous to the dentate nuclei in mammals. In a first group of birds (n=7), a unilateral electrolytic lesion was performed in the lateral deep cerebellar nucleus by passing 0.05mA during 30 seconds through a tungsten electrode. Lesions were made at three points (see the stereotaxic coordinates in Table1, DCN coordinates, second row). In a second experimental group (n=3), chemical partial lesion was performed using ibotenic acid in 1µL Hamilton syringe, with a rate of 100nL/min. We also performed injections at three locations (see Table1, DCN coordinates) injecting 150nL per point. Sham lesions were performed in another group of age-matched juvenile birds. Sham birds underwent the same surgery as the lesion group, with a stimulating electrode was placed at the lesion location but no current was applied. Both lesion and sham protocols were done around 57 days post hatch (56,8 +/- 7,5 days post hatch for lesion group, 57.0 +/- 4,5 days post hatch for sham group). Following surgery, the behavior of birds was closely monitored for a few days to ensure proper recovery. Many birds underwent temporary motor deficits (postural and balance troubles) for a couple of days but recovered very quickly and were all perching and feeding normally 48h after surgery. Singing usually resumed after 48h, or at most after 72h. Each juvenile (sham and lesion) was put in a recording box one week before the lesion experiment, and recorded using Sound Analysis Pro software (SAP, Tchernichovski et al., 2001). To prevent any deficit due to the lack of tutor, we presented the tutor to the juvenile two hours per day until the bird underwent the surgery. All birds had same access to their respective tutors. After the surgery, each juvenile was recorded until the crystallization phase (30 days after the surgery experiment).

### - Histology

For the anatomical tracing protocol: Birds were sacrificed with a lethal intraperitoneal injection of pentobarbital (Nembutal, 54.7mg/mL), perfused intracardially with PBS 0.01M followed by 4% paraformaldehyde as fixative. The brain was removed, post-fixed in 4% for 24h, and cryoprotected in 30% sucrose. We then cut 40µm thick sections in the parasagittal plane with a freezing microtome. Slices were mounted with Mowiol (Sigma Aldrich) and observed under an epifluorescence (Leica Microsystems, Leica DM 1000, Nanterre, France) or a confocal microscope (Zeiss, LSM 710, France). Images were analyzed using ImageJ software (Rasband WS, NIH, Bethesda, Maryland, USA).

After electrophysiological recordings, the bird was perfused as described above. Then, brain was removed, post-fixed one day in PFA 4%, store in sucrose 30%, and we did 60µm slices with Nissl staining to control the stimulation electrode and recording electrode tracts.

For the lesion protocol: All juvenile birds were sacrificed at 100 dph using the protocol previously described for tracing protocol. We then cut 60µm-thick cerebellar sections in the horizontal plane with a freezing microtome. We did Nissl staining to check lesions locations. Slices were mounted with Mowiol (Sigma Aldrich) and observed under a transmitted-light microscope (Leica Microsystems, Leica DM1000, Nanterre, France). With ImageJ software (Rasband WS, NIH, Bethesda, Maryland, USA), we calculated the area of lesion for each nucleus compared to the control nucleus in the other hemisphere.

### - Song analysis

Songs were continuously recorded using Sound Analysis Pro software (SAP, Tchernichovski et al., 2001). Songs were then sorted and analyzed using custom Matlab (MathWorks, Natick, MA, USA) programs. Briefly, the program detected putative motifs based on peaks in the cross-correlation between the sound envelope of the recorded sound file and a clean preselected motif. Putative motifs were then sorted based on their spectral similarity with the pre-selected clean motif, using thresholds set by the experimenter. This analysis allowed us to successfully sort >98% of the songs produced by a bird on a given day (assessed by comparing hand sorting with the automated sorting by the program). We calculated the spectrogram of each extracted song (fast Fourier transforms using 256-point Hanning windows moved in 128-point steps).

For each family including a juvenile bird undergoing DCN lesion or sham-lesion, the spectrograms of 10 randomly-selected and manually checked renditions of the stereotyped motif produced by the tutor were stored for comparison with the juvenile’s songs. Among all songs produced by the juvenile in each considered condition: before lesion or at crystallization (all recordings from a single day of recording were considered for analysis in each condition: pre-surgery or after crystallization), 10 randomly-selected songs were compared to the tutor’s selected motifs using the following procedure. Cross-correlations were computed between all possible pairs of this subset. For each pair consisting of a tutor’s motif and a juvenile’s song, a cross-correlation index was calculated as the sum of the cross-correlation function between their two spectrograms, normalized by the square root of the product of their auto-correlation function. The average cross-correlation index over all 100 pairs was called the ‘spectral similarity index’ between tutor and juvenile in that condition.

### - Statistics

Numerical values are given as mean ± SD, unless stated otherwise.

Electrophysiology: As the goal of pharmacological experiments was to look at the effect of glutamatergic transmission blockade on baseline response strength induced by DCN stimulation, we compared the mean response strength during two conditions: the baseline condition and the drug condition. To do so we performed a paired Wilcoxon test between the control response and that after application of drugs. We used non-parametric statistical tests because of the small number of neurons recorded (less than 30 neurons in each experiment).

Behavior: Given our initial hypothesis that the cerebellum may contribute to song learning, we planned to compare the similarity between juvenile and tutor songs before and after surgery, as well as at crystallization (90 dph). This comparison was applied both in sham-lesion birds and in DCN lesion birds. The similarity scores in these two groups were compared between pre-surgery and crystallization period using a paired Wilcoxon test (MathWorks, Natick, MA, USA). Additionally, we tested whether there was a significant correlation between the size of the lesion and the improvement in tutor song imitation after surgery. To this end, we calculated the correlation coefficient between the lesion size (proportion of DCN left unaffected, determined histologically for DCN lesion birds, and assigned to 100% for sham-lesion birds) and the normalized song similarity at crystallization (similarity at 90 days post hatch / similarity before surgery). We tested the hypothesis of no correlation: each p-value was determined as the probability of obtaining a correlation larger than the observed value by chance, when the true correlation is zero (MathWorks, Natick, MA, USA).

## Acknowledgements

We are grateful to Carole Levenes for valuable discussions and to Claude Meunier, David Hansel and David J Perkel for their comments on the manuscript. This work was supported by the Agence National pour la Recherche (ANR, program “Retour Post-Doc”, Grant number ANR-10-PDOC-0016) and by the city of Paris (program “Emergence”, Grant number DDEEES 2014-166).

## Competing interest

No competing interests declared.

